# A sterol-binding pocket in iRhom1 underlies paralog-specific regulation of the sheddase ADAM17

**DOI:** 10.64898/2026.04.28.721421

**Authors:** Fangfang Lu, Hongtu Zhao, Yaxin Dai, Chia-Hsueh Lee, Matthew Freeman

## Abstract

ADAM17, the major sheddase in mammalian cells, releases membrane-tethered EGFR ligands and inflammatory cytokines, and is a central regulator of cell signalling. The rhomboid pseudoproteases, iRhom1 and iRhom2, function as essential cofactors of ADAM17, controlling its maturation and activation. In contrast to the well-characterized iRhom2, the mechanism and regulation of its ubiquitously expressed paralog iRhom1 remain undefined. Here, we present a 2.5 Å cryo-EM structure of the full-length human iRhom1/ADAM17 complex, revealing a previously unrecognized sterol-binding pocket located between TMD2 and TMD5. Structure-guided mutagenesis and pharmacological perturbation of sterol binding demonstrate that sterol binding is required to stabilize the iRhom1/ADAM17 complex and sustain its shedding activity. Strikingly, this regulation is paralog-specific: iRhom2 precludes sterol binding and instead stabilizes ADAM17 through direct intramolecular interactions. Furthermore, two human iRhom1 variants associated with cardiac disease localize adjacent to the sterol-binding pocket and disrupt ADAM17 maturation and activity. Together, these findings uncover mechanistic divergence between iRhom paralogs and establish a sterol-binding pocket in iRhom1 as a critical determinant of ADAM17 stability, revealing a potential avenue for paralog-selective therapeutic targeting.

## Introduction

The metalloprotease ADAM17 is the principal sheddase in mammalian cells, responsible for the proteolytic release of diverse membrane-tethered substrates, including tumor necrosis factor (TNF) and multiple epidermal growth factor receptor (EGFR) ligands ^1^. Consistent with its central role in signalling, dysregulated ADAM17 is associated with a wide variety of diseases, including inflammatory skin and bowel disease ^2,3^, chronic kidney diseases ^4^, cardiovascular diseases ^5^, Alzheimer’s disease ^6^, as well as multiple cancers ^7,8^. Despite its broad clinical relevance, the molecular mechanisms that govern ADAM17 maturation and stability remain incompletely understood.

ADAM17 function critically depends on the inactive rhomboid-like pseudoproteases iRhom1 and iRhom2, which form a 1:1 complex with the protease in cellular membranes. iRhoms regulate ADAM17 at multiple levels. First, iRhoms are essential for the release of immature ADAM17 from the endoplasmic reticulum (ER), thereby allowing its trafficking to the Golgi apparatus and beyond, where it is matured by release of its inhibitory prodomain by furin ^9,10^. Second, at the plasma membrane, iRhoms regulate ADAM17 activation: cellular stimulation via GPCR signalling leads to phosphorylation of the iRhom2 cytoplasmic domain, thereby recruiting 14-3-3 proteins to trigger ADAM17 activation ^11,12^. Recent structural work on the iRhom2/ADAM17 complex revealed how iRhom2 restrains ADAM17 activity and stabilizes the mature protease, providing important mechanistic insight into sheddase regulation ^13,14^.

Both iRhom1 and iRhom2 support ADAM17 maturation and have generally been assumed to function through similar molecular mechanisms. However, evidence suggests they may not be fully redundant. The two paralogs exhibit distinct expression patterns: iRhom2 is enriched in immune cells and skin, whereas iRhom1 is more ubiquitously expressed ^15^. Consistent with these differences, knockout of iRhom1 in mice results in early lethality with phenotypes including brain haemorrhage and cardiac defects, whereas iRhom2-deficient mice are viable, displaying predominantly inflammatory and immune defects when challenged ^15,16^. Differences in substrate selectivity and stress-induced expression have also been reported ^17–19^. However, in the absence of structural information for iRhom1, whether the two paralogs regulate ADAM17 through mechanistically distinct principles remains unresolved.

Here, we report the cryo-EM structure of the full-length human iRhom1/ADAM17 complex, challenging the prevailing view that the two paralogs operate through equivalent mechanisms. Unexpectedly, we identify a sterol-binding pocket within the conserved rhomboid-like core of iRhom1, positioned between transmembrane domains (TMD) 2 and 5 at a site corresponding to the substrate entry cavity of active rhomboid proteases. Functional analyses demonstrate that this sterol-binding pocket is required for stable formation of the mature iRhom1/ADAM17 complex. Disruption of sterol engagement destabilizes mature ADAM17, impairs its trafficking to the cell surface, and reduces shedding activity. Strikingly, this mechanism is specific to iRhom1: the equivalent region in iRhom2 adopts a closed conformation and does not exhibit sterol-dependent stabilization. Together, our findings uncover a previously unrecognized structural determinant of ADAM17 stability and reveal fundamental mechanistic divergence between iRhom paralogs.

## Results

### Cryo-EM structure of the human iRhom1/ADAM17 complex reveals a sterol-binding pocket in iRhom1

To study the structure of the human iRhom1/ADAM17 complex, we co-expressed HEK cells full-length ADAM17, iRhom1 and the accessory protein FRMD8, which stabilises the complex ^12,20^. Using single-particle cryo-electron microscopy (cryo-EM), we determined the structure of the complex at 2.5 Å resolution (Figures 1 and S1). As previously observed for the iRhom2/ADAM17 complex ^13^, the cytoplasmic domain of iRhom1, including bound FRMD8, was not resolved due to conformational flexibility. Nevertheless, the high-resolution density map allowed us to unambiguously model the extracellular and transmembrane regions of iRhom1 and ADAM17 (Figure 1A, B).

**Figure 1.**
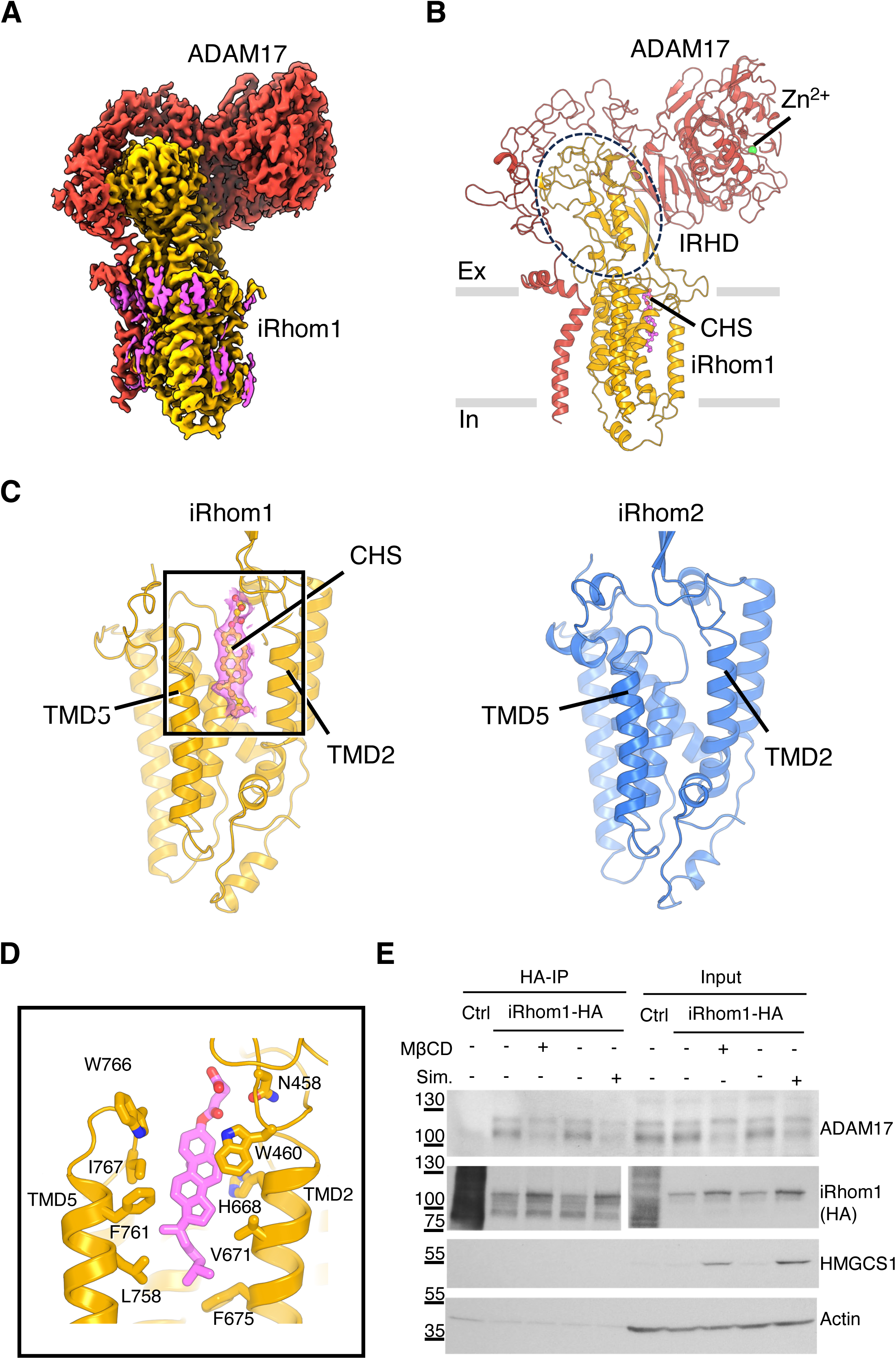
Cryo-EM structure of the human iRhom1/ADAM17 complex reveals a sterol-binding pocket in iRhom1. (A) Cryo-EM density map of the human iRhom1/ADAM17 complex. ADAM17 is shown in red, iRhom1 in yellow, and the bound lipid density in magenta. (B) Structural model of the iRhom1/ADAM17 complex shown in cartoon representation. Adam17 in red, iRhom2 in yellow, the zinc ion in green. The iRhom homology domain (IRHD) is highlighted in dashed circle line, and the modelled CHS molecule is shown in magenta. Ex: extracellular; In: intracellular. (C) Comparison of the transmembrane domains of iRhom1 and iRhom2. iRhom1 contains a sterol-binding pocket between TMD2 and TMD5 occupied by CHS. No similar density is found in iRhom2. (D) Close-up view of the iRhom1 sterol-binding pocket. Residues lining or adjacent to the CHS-binding site are labelled. (E) Cholesterol depletion reduces the association between iRhom1 and mature ADAM17. HT-29 cells stably expressing HA-tagged iRhom1 were treated with methyl-β-cyclodextrin (5mM MβCD, 3 h) or simvastatin (2 µM, 48 h). Cells stabling expressing HA-tagged UNC93B served as control (Ctrl). HA-based immunoprecipitation was performed and the associated ADAM17 was detected by immunoblotting. Input lysates were blotted for ADAM17, iRhom1-HA, and actin. Increased HMGCS1 level indicates successful cholesterol depletion.

The complex comprises one molecule each of iRhom1 and ADAM17, with the two proteins interacting extensively in both the extracellular and transmembrane regions (Figure 1A, B). The extracellular domain of iRhom1 (the iRhom homology domain, IRHD) extends approximately 45 Å above the membrane. This domain serves as a molecular signature of iRhom proteins and is not found in other rhomboid proteases or pseudoproteases. The transmembrane region of iRhom1 adopts the conserved rhomboid-like fold, reminiscent of the helical arrangement observed in other rhomboid family members. Overall, the architecture of the iRhom1/ADAM17 complex closely resembles that of the previously reported iRhom2/ADAM17 structure.

Notably, however, we observed a well-defined density located between transmembrane domains (TMDs) 2 and 5 of iRhom1 that is absent in the iRhom2 structure (Figure 1C). Inspection of this density revealed features of a cholesterol-like molecule and, as cholesteryl hemisuccinate (CHS) was present during protein purification, we modelled the density as a CHS molecule. The insertion of CHS led to TMD2 and TMD5 of iRhom1 being more distant from each other than they are in iRhom2 (Figure S1F), in turn causing a shift of the position of the IRHD of iRhom1 by ∼5 Å, and a consequent 6 Å displacement of the ADAM17 extracellular domain (Figure S1G).

The location of sterol binding in iRhom1 is noteworthy. In characterised rhomboid proteases and the rhomboid pseudoprotease Derlin, the space between TMD2 and TMD5 is the site where substrate or client TMDs bind, facilitating entry and docking ^21,22^. In iRhom1, this protein substrate/client binding site has been repurposed as a sterol-binding pocket, lined by hydrophobic residues, including V671 and F675 of TMD2, as well as L758, F761, I767, and W766 of TMD5 (Figure 1D). Additionally, H668 of TMD2 and N458 and W460 of IRHD are also nearby. Conservation analysis of iRhom1 across species showed that these sterol-binding residues are highly conserved (Figure S2A, B), suggesting their functional importance.

### Sterol engagement contributes to stability of the iRhom1/ADAM17 complex

To examine the functional consequences of sterol occupancy in iRhom1, we first assessed the effect of cholesterol depletion on the iRhom1/ADAM17 complex. Human HT-29 cells stably expressing HA-tagged iRhom1 were treated with either simvastatin (48 h; chronic depletion) or methyl-β-cyclodextrin (MβCD; 3 h; acute depletion). Under both conditions, the interaction between iRhom1 and mature ADAM17 was markedly reduced (Figure 1E), indicating that membrane cholesterol contributes to stabilization of the iRhom1/mature ADAM17 complex.

We next examined the functional importance of residues lining the sterol-binding pocket. In iRhom1/iRhom2 double-knockout (DKO) cells, defective ER exit means that only the immature form of ADAM17 is present (Figure 2A). Expression of wild type (WT) iRhom1 restored robust ADAM17 maturation, as evidenced by accumulation of the lower molecular weight mature form. In contrast, mutating predicted sterol-interacting residues (N458A/W460A, H668A, F675A) substantially reduced mature ADAM17 levels; protein expression levels of the mutants were comparable to WT (Figure 2A). Two additional neighbouring residues (W766A/I767A) had minimal effect. These findings indicate that integrity of the sterol-binding pocket is important for accumulation of mature ADAM17.

**Figure 2.**
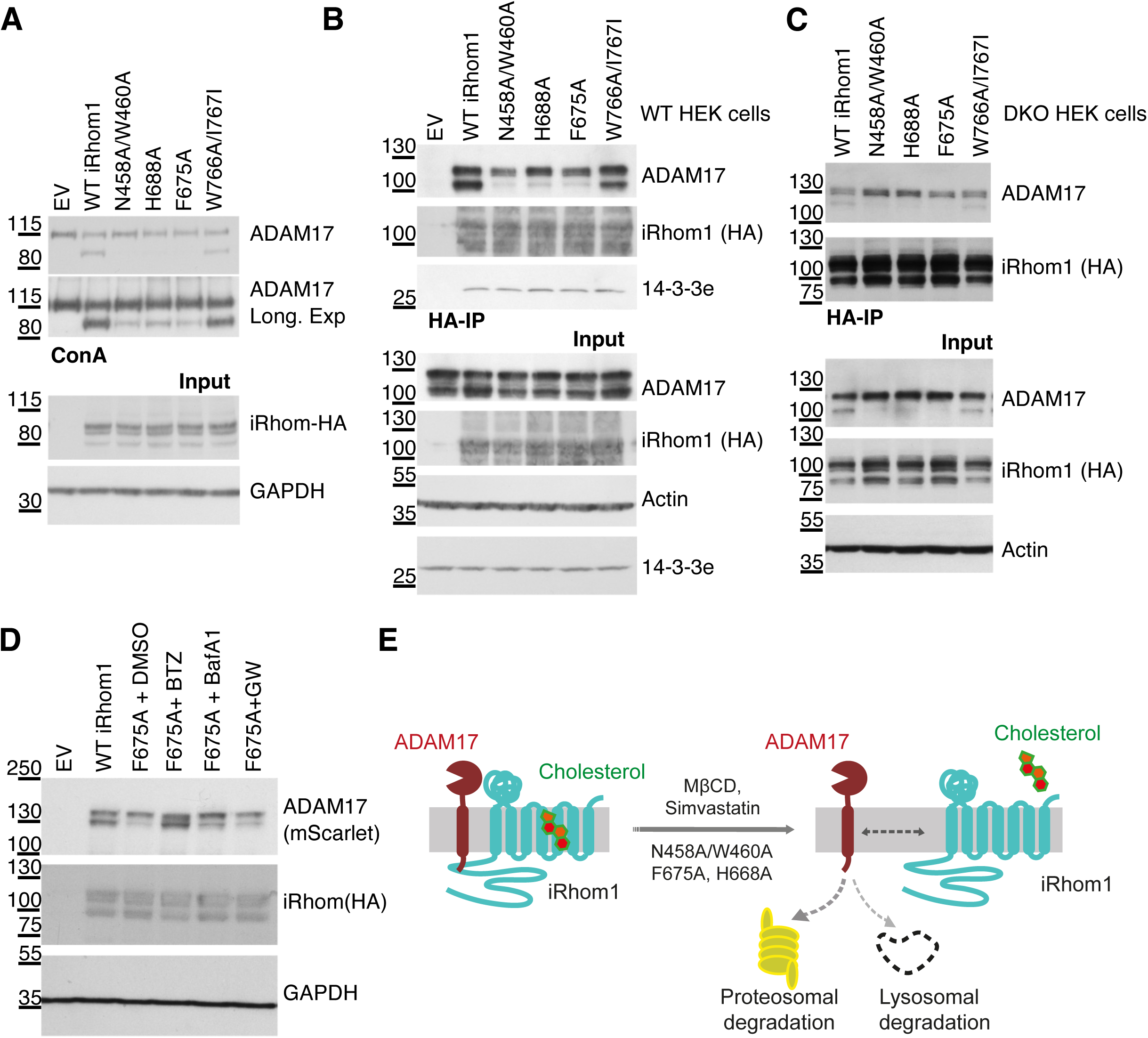
Integrity of the sterol-binding pocket is required for stabilization of mature ADAM17 by iRhom1. (A) iRhom1/2 double-knockout HEK293T cells were transfected with empty vector, wild-type iRhom1, or the indicated iRhom1 sterol-pocket mutants. ADAM17 maturation was assessed by ConA enrichment and immunoblotting. Input lysates were analysed for iRhom1-HA and GAPDH. (B) Wild-type HEK293 cells expressing the indicated iRhom1 variants were subjected to HA immunoprecipitation. Co-precipitated ADAM17, iRhom1-HA and 14-3-3ε were detected by immunoblotting. Input lysates were analysed for ADAM17, iRhom1-HA, actin and 14-3-3ε. The binding to important signalling protein 14-3-3ε was not affected by the iRhom1 mutants. (C) iRhom1/2 double-knockout HEK293T cells were transfected with indicated iRhom1 variants and HA immunoprecipitation was performed to assess iRhom binding to ADAM17. (D) Western blots of iRhom1/2 DKO HEK293 cells transfected with iRhom1 mutants together with wild-type (WT) mScarlet-tagged ADAM17. Cells were treated with DMSO, the proteasome inhibitor bortezomib (BTZ, 1 μM), the lysosomal inhibitor bafilomycin A1 (BafA1, 1 μM), or the ADAM17 inhibitor GW280264X (2 μM) for 12 h before harvesting. Mature ADAM17 and iRhom1-HA levels were analysed by immunoblotting. (E) Schematic showing sterol-dependent stabilization of the iRhom1/ADAM17 complex. Disruption of sterol binding by cholesterol depletion or sterol-pocket mutation destabilizes mature ADAM17 and promotes its degradation.

Because mutations in residues in the sterol-binding pocket reduced mature ADAM17 levels, we next tested whether these mutations affected the ability of iRhom1 to bind ADAM17. In wild-type HEK293 cells, which contain both immature and mature ADAM17, mutations in the sterol-binding pocket (N458A/W460A, H668A, F675A, but not the W766A/I767A) selectively reduced association with mature ADAM17, while interaction with the immature form was largely preserved (Figure 2B). Similar results were obtained in DKO cells (Figure 2C). Together, these data suggest that residues predicted to coordinate sterol engagement are required to stabilize interaction specifically with mature ADAM17.

The reduction in mature ADAM17 could reflect either impaired maturation or decreased stability of the mature form. Because interaction between iRhom1 and immature ADAM17 was unaffected, we believe that a primary defect in ER exit or early trafficking is unlikely. We therefore examined ADAM17 stability. In cells expressing sterol-pocket mutants, mature ADAM17 levels were markedly reduced (Figure 2D). This reduction was rescued by the proteasome inhibitor bortezomib and partially restored by the lysosomal inhibitor bafilomycin A1, indicating enhanced degradation of mature ADAM17 when bound to the mutant forms of iRhom1. In contrast, inhibition of ADAM17 catalytic activity with GW280264X did not restore protein levels, suggesting that degradation is not secondary to protease activation. Together, these findings support a model in which sterol binding to iRhom1 contributes to stabilization of the mature iRhom1/ADAM17 complex (see illustration in Figure 2E).

### Sterol-dependent stabilization of the ADAM17 complex is specific to iRhom1

Although iRhom1 and iRhom2 have generally been believed to have the same function in ADAM17 regulation, our structural data revealed a clear distinction: under identical purification conditions, only the iRhom1/ADAM17 complex accommodated a sterol molecule at the TMD2-TMD5 interface; no comparable density was observed in the iRhom2 structure (Figure 1C). This observation suggests that sterol-dependent stabilization may be specific to iRhom1, which was confirmed by showing that, in HT-29 cells, cholesterol depletion markedly disrupted the interaction between iRhom1 and mature ADAM17 but had minimal effect on the association between mature ADAM17 and either of the two iRhom2 isoforms (Figure 3A).

**Figure 3.**
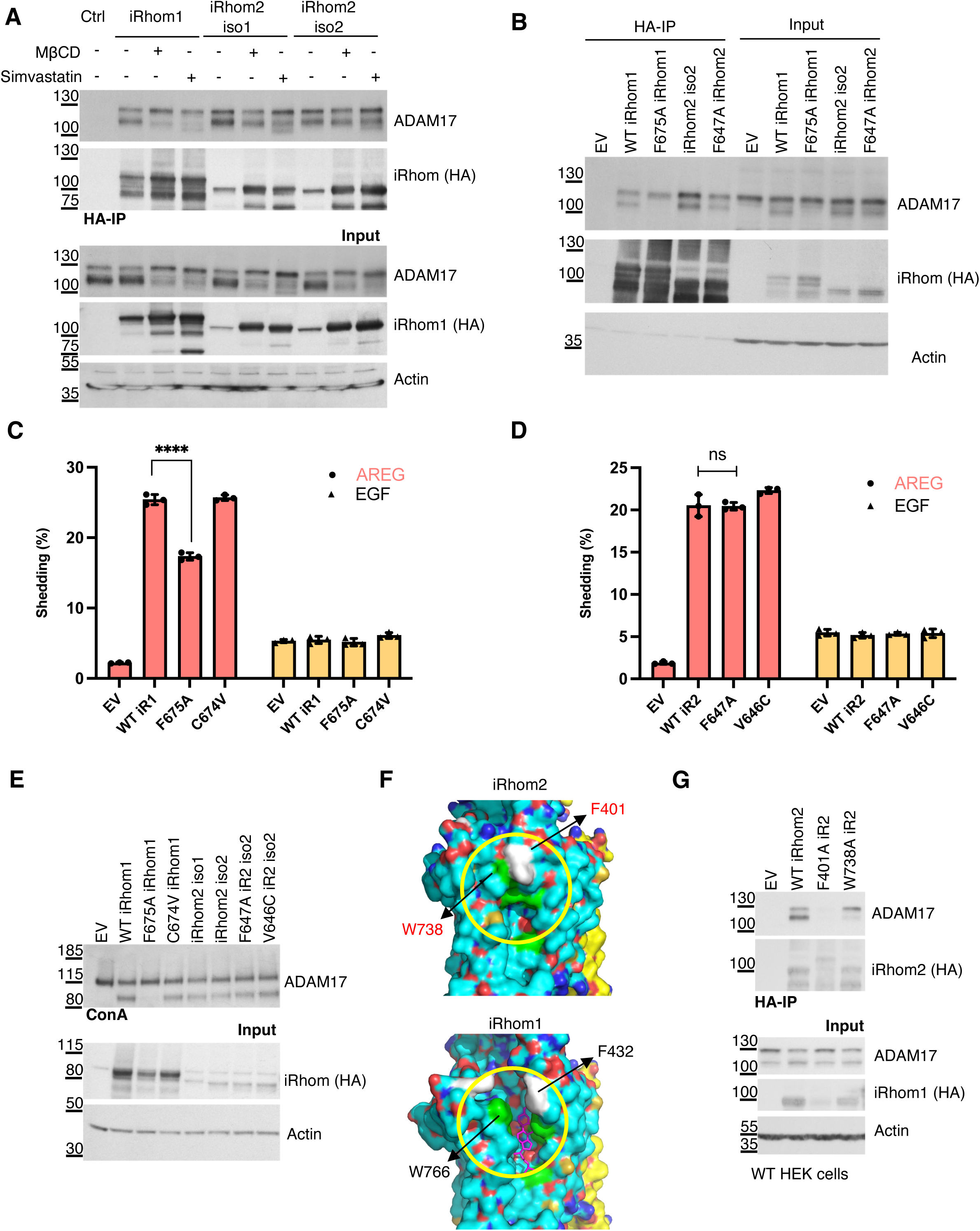
Sterol-dependent stabilization of ADAM17 is specific to iRhom1. (A) HT-29 cells stably expressing HA-tagged iRhom1 or iRhom2 isoforms were treated with MβCD or simvastatin, as described above. HA immunoprecipitation was used to assess the interaction between iRhom proteins and ADAM17. (B) HA immunoprecipitation analysis of iRhom1/2 double-knockout HEK293T cells re-expressing wild-type or mutant iRhom1 and iRhom2 variants. Co-precipitated ADAM17 and iRhom-HA were detected by immunoblotting. (C, D) ADAM17-dependent shedding assays in cells expressing wild-type or mutant iRhom1 (C) or iRhom2 (D). Shedding of alkaline phosphatase-tagged AREG and EGF substrates was quantified. Data are shown as mean ± standard deviation (SD; n = 3, three transfectants); ns = not significant, **** = p < 0.0001. Data from three independent experiments were collated in Figure S3. (E) ConA enrichment analysis of ADAM17 maturation in iRhom1/2 double-knockout HEK293T cells re-expressing wild-type or mutant iRhom1 and iRhom2 variants. (F) Structural comparison of the TMD2-TMD5 region in iRhom1 and iRhom2. iRhom1 adopts an open conformation that accommodates sterol binding, whereas iRhom2 adopts a closed conformation stabilized by residues including F401 and W738. (G) HA immunoprecipitation analysis showing that mutation of iRhom2 residues F401 or W738 disrupts association with ADAM17, consistent with a direct IRHD-TMD core stabilization mechanism in iRhom2.

We next assessed any potential function for iRhom2 residues corresponding to those in the iRhom1 sterol-binding pocket. Although pocket residues are conserved between the two paralogs, an F647A mutation in iRhom2 (corresponding to F675A in iRhom1) did not significantly affect ADAM17 interaction or maturation (Figure 3B, E), nor did it alter PMA-stimulated shedding activity (Figure 3C, D). In contrast, the corresponding iRhom1 F675A mutation substantially reduced mature ADAM17 levels and impaired shedding. These results indicate that the TMD2-TMD5 interface contributes differently to ADAM17 regulation in the two paralogs.

Given the conservation of sterol-contacting residues, why does iRhom2 not bind cholesterol? We asked whether neighbouring residues are responsible for the difference. Sequence analysis identified adjacent residues that differ between the paralogs (C674 in iRhom1 and V646 in iRhom2) near the conserved phenylalanine (F675/F647). However, reciprocal mutation of residues adjacent to F675/F647 that differ between the paralogs (C674V in iRhom1 and V646C in iRhom2) did not affect ADAM17 maturation or shedding (Figure 3C-E), indicating that simple local sequence differences do not account for paralog-specific cholesterol binding. We therefore turned to structural comparison. Notably, iRhom1 adopts an open conformation at the TMD2-TMD5 interface that can accommodate sterol binding (Figure 3F, upper panel), whereas iRhom2 maintains a closed conformation at the equivalent region (Figure 3F, lower panel), restricting sterol access. In iRhom2, this closed state appears stabilized by a π-stacking interaction between F401 in the IRHD and W738 in TMD5, anchoring the IRHD to the rhomboid core. Disruption of this interaction by mutating F401A or W738A significantly reduced iRhom2-ADAM17 binding (Figure 3G), demonstrating that direct IRHD-core contacts contribute to complex stability in iRhom2. The corresponding mutation W766A in iRhom1 (in combination with I767A) had minimal effect on the iRhom1/ADAM17 complex (Figure 2A-C).

Overall, given the position of the sterol-binding pocket within the rhomboid core and the involvement of IRHD residues (N458, W460) in stabilizing the iRhom1/ADAM17 complex, we propose that sterol engagement in iRhom1 may act as a molecular bridge between the IRHD and transmembrane core. In contrast, iRhom2 achieves comparable stabilization through direct π-stacking interactions between IRHD and TMD5. Thus, although both paralogs rely on correct IRHD positioning to maintain ADAM17 binding, they use distinct structural strategies to achieve that conformation.

### Sterol-pocket mutations reduce cell-surface abundance of the iRhom1/ADAM17 complex

Having established that sterol-binding stabilizes the iRhom1/mature ADAM17 interaction and thereby protects mature ADAM17 from degradation, we next examined whether disruption of the sterol-binding pocket affects cell-surface abundance of the complex. Mutations in iRhom1 residues that we found to participate in sterol binding (N458A/W460A, H668A, F675A) significantly reduced cell-surface levels of both iRhom1 and ADAM17 compared to wild-type (WT) controls (Figure 4A-D). In contrast, the W766A/I767A mutant, which had minimal effects on complex stability and ADAM17 maturation in previous assays, exhibited near-normal surface expression. This indicates that integrity of the sterol-binding pocket is required not only to stabilise of the mature ADAM17 complex but also to maintaining normal levels of iRhom1 at the plasma membrane.

**Figure 4.**
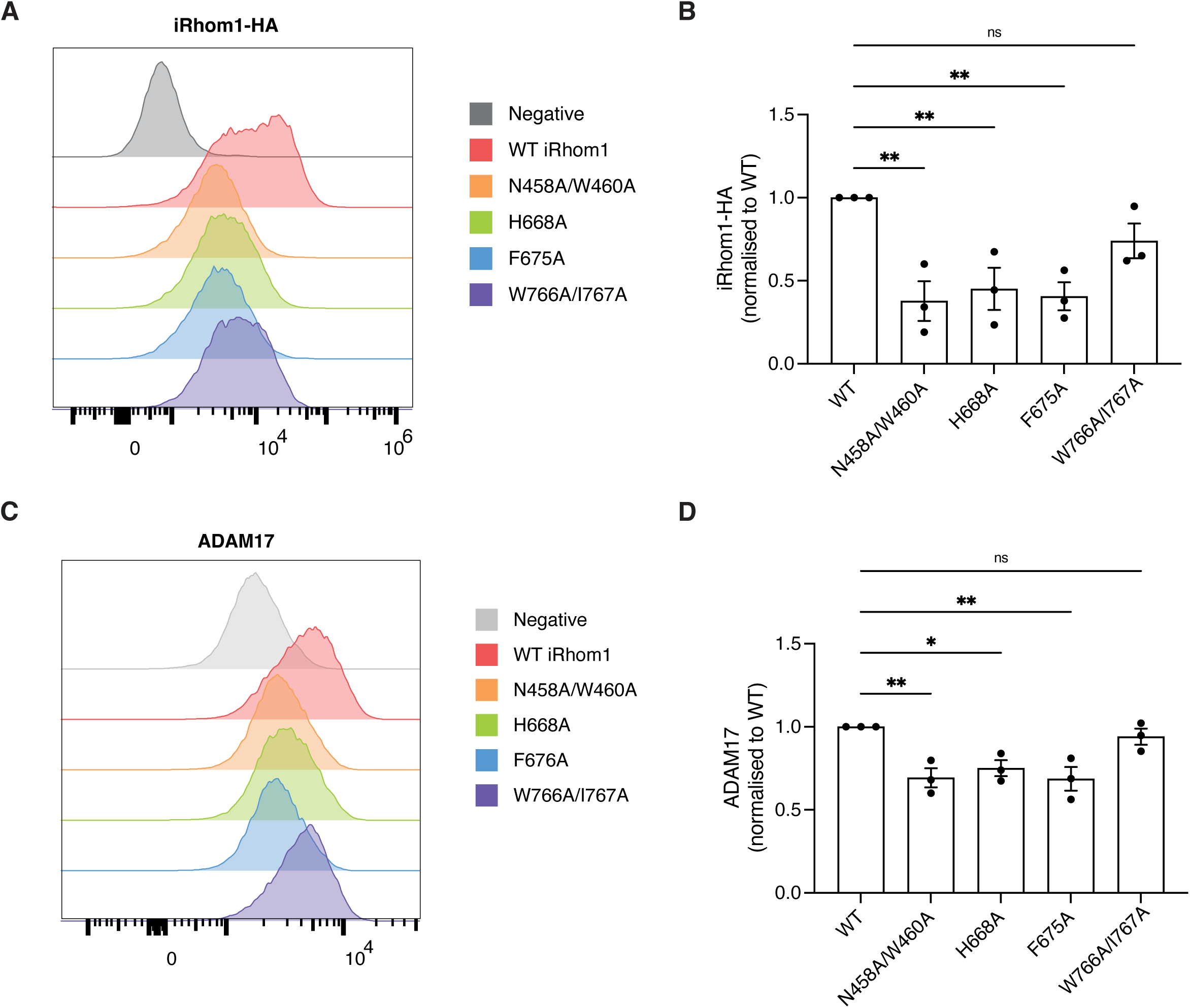
Sterol-pocket mutations reduce cell-surface abundance of iRhom1 and ADAM17. (A) Representative flow cytometry histograms showing cell-surface levels of HA-tagged iRhom1 variants expressed in iRhom1/2 double-knockout HEK293T cells. (B) Quantification of cell-surface iRhom1-HA levels for various mutants, normalized to wild-type iRhom1. Data are from three independent experiments. (C) Representative flow cytometry histograms showing cell-surface ADAM17 levels in DKO cells expressing wild-type or mutant iRhom1. (D) Quantification of cell-surface ADAM17 levels, normalized to cells expressing wild-type iRhom1. Data are shown as mean ± standard error of the mean (SEM); ns = not significant, * = p < 0.05, ** = p < 0.01, **** = p < 0.0001.

### Disease-associated iRhom1 mutations cluster near the sterol-binding pocket and disrupt ADAM17 function

Mutations in the cytoplasmic domain of iRhom2 are known to cause the rare cancer syndrome tylosis with oesophageal cancer (TOC) ^23,24^. In contrast, no iRhom1 variants have been linked to this syndrome. Instead, rare missense mutations in *RHBDF1* (encoding iRhom1) have been associated with cardiac pathologies, including Wolff-Parkinson-White (WPW) syndrome and Childhood-Onset Cardiomyopathy ^25,26^ (Figure 5A). In both cases, homozygous mutations (G462R in WPW syndrome and G665W in cardiomyopathy) were identified in affected families, and the genetic data suggest a causal role, although the underlying molecular mechanisms remain unexplored.

**Figure 5.**
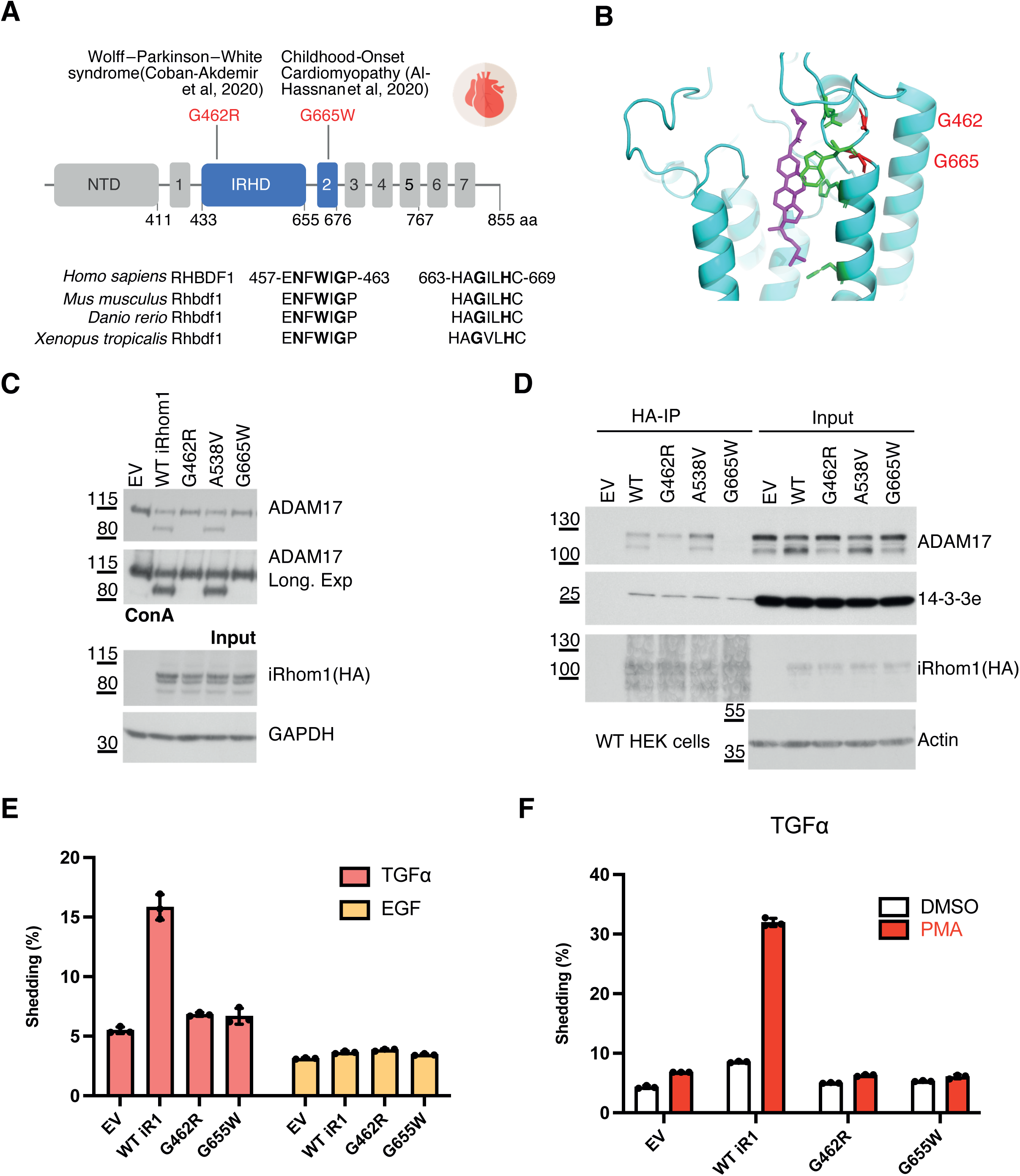
Disease-associated iRhom1 variants near the sterol-binding pocket impair ADAM17 maturation and shedding. (A) Schematic of human iRhom1 protein showing the positions of the disease-associated variants G462R and G665W. G462R has been associated with Wolff–Parkinson–White syndrome, and G665W with childhood-onset cardiomyopathy. Conservation of the surrounding regions across vertebrate iRhom1 orthologues is shown below. NTD: N-tereminal domain. Transmembrane domains are numbered. (B) Structural location of G462 and G665 (in red) relative to the sterol-binding pocket in iRhom1. The CHS molecule is shown in magenta. (C) ADAM17 maturation was assayed by ConA enrichment and immunoblotting using iRhom1/2 double-knockout HEK293T cells re-expressing wild-type iRhom1, disease-associated iRhom1 variants, or the control variant A538V. (D) HA immunoprecipitation analysis of wild-type HEK293 cells expressing the indicated iRhom1 variants. Co-precipitated ADAM17 and 14-3-3ε were detected by immunoblotting. The binding to important signalling protein 14-3-3ε was not affected by the iRhom1 mutants tested. (E) Constitutive ADAM17-dependent shedding of alkaline phosphatase-tagged TGFα and EGF substrates in DKO cells re-expressing wild-type or mutant iRhom1. (F) PMA-stimulated shedding of alkaline phosphatase-tagged TGFα in cells expressing wild-type or mutant iRhom1. Data are shown as mean ± SD (n=3, three transfectants).

Intriguingly, both disease-associated mutations localize adjacent to the sterol-binding pocket identified in our structure. G462R lies near N458 and W460 within the IRHD, whereas G665W is positioned close to H668 in TMD2 (Figure 5A, B). These residues are highly conserved across vertebrates, underscoring their likely functional importance. We examined whether these mutations support ADAM17 maturation and activity. When expressed in iRhom1/2 double-knockout (DKO) cells, both G462R and G665W mutants severely impaired ADAM17 maturation (Figure 5C) and abolished ADAM17-dependent shedding, both constitutively (Figure 5E), and following PMA stimulation (Figure 5F). Interestingly, co-immunoprecipitation analysis revealed mechanistic differences between the mutants: G462R specifically disrupted interaction with mature ADAM17, whereas G665W abolished interaction with both immature and mature ADAM17 (Figure 5D; confirmed in DKO cells, Figure S3C). These findings indicate that both variants, which are located adjacent to the sterol binding pocket in iRhom1, compromise stability of the iRhom1/ADAM17 complex, albeit through distinct mechanisms. As a specificity control, a nearby benign variant A538V had no detectable effect on ADAM17 maturation or shedding.

## Discussion

ADAM17 is an essential sheddase in mammalian cells, regulating the release of cytokines, growth factors, and adhesion molecules that control inflammatory, cardiovascular, and oncogenic signalling pathways ^5,27,28^. Despite its clinical importance, direct therapeutic targeting of ADAM17 has been largely unsuccessful, in part because its regulatory mechanisms remain incompletely understood. Structural resolution of the iRhom2/ADAM17 complex provided important insight into how the pseudoprotease iRhom2 stabilizes and restrains ADAM17 activity ^13,14^.

However, the molecular mechanisms governing regulation by the ubiquitously expressed paralog iRhom1 have remained unclear. Here, we identify a sterol-binding pocket within the rhomboid core of iRhom1 that is structurally and functionally required for stabilization of the mature iRhom1/ADAM17 complex. Disruption of sterol engagement, either by cholesterol depletion or mutation of pocket residues, destabilizes mature ADAM17, leading to enhanced degradation and reduced surface abundance. These findings reveal a previously unrecognized sterol-dependent mechanism that contributes to maintaining ADAM17 complex integrity.

A conceptual advance of this study is the demonstration that the two iRhom paralogs stabilize ADAM17 through distinct structural mechanisms. Our previous work established that proper interaction between the iRhom2 IRHD and the ADAM17 ectodomain is essential for maintaining the mature complex ^13^. In the present study, we show that both paralogs require stabilization of the IRHD to preserve productive interaction with ADAM17 but achieve this through different mechanisms. In iRhom2, direct π-stacking between F401 and W738 anchors the IRHD to the rhomboid core. In contrast, iRhom1 accommodates a sterol molecule at the TMD2-TMD5 interface, which likely acts as a molecular bridge between the IRHD and the transmembrane core. Thus, although the overall architecture of the two complexes is highly similar, they have evolved paralog-specific means of maintaining IRHD-dependent stabilization of the sheddase complex.

Our findings are consistent with emerging evidence that membrane lipids can function as structural cofactors in multi-pass membrane proteins. For example, cholesterol binding within the transmembrane domain of tetraspanin CD81 induces conformational changes that regulate its interaction with client proteins ^29^. Similarly, prior in vivo studies have reported that cholesterol depletion via simvastatin treatment reduces ADAM17 protein levels in mice ^30^. Although the mechanism underlying that observation was unclear, our results raise the possibility that cholesterol-dependent stabilization of the iRhom1/ADAM17 complex may contribute to this effect. Together, these observations suggest that lipid occupancy within transmembrane domains may represent a broader regulatory principle in membrane protein complexes.

The location of the sterol-binding pocket between TMD2 and TMD5 is particularly intriguing. This region corresponds to the canonical substrate-entry site in catalytically active rhomboid proteases ^22^ and has been proposed as a potential interaction site for client transmembrane segments ^17^. The presence of a sterol molecule within this cavity raises the possibility that lipid occupancy may influence conformational dynamics or substrate accessibility in an iRhom1-specific manner. Further structural and computational studies incorporating sterol molecules will be required to explore this possibility.

Importantly, two human iRhom1 variants associated with cardiac disease (G462R and G665W) localize adjacent to the sterol-binding pocket and profoundly disrupt ADAM17 maturation and activity. These findings provide a mechanistic link between RHBDF1 mutations and defective ADAM17 regulation. Although direct biochemical measurements of sterol binding will be necessary to establish causality, the clustering of pathogenic variants near the sterol-proximal structural region underscores its functional importance.

Collectively, our study reveals sterol engagement as a previously unrecognized determinant of iRhom1-mediated ADAM17 stabilization and uncovers fundamental mechanistic divergence between iRhom paralogs. The presence of a sterol-accessible cavity within iRhom1 suggests the potential for selective pharmacological modulation of the iRhom1/ADAM17 complex, highlighting a potential avenue for paralog-selective modulation of ADAM17 activity.

## Methods and Materials

### Key resources table

| REAGENT or RESOURCE | SOURCE | IDENTIFIER |
| --- | --- | --- |
| **Antibodies** |  |  |
| Rabbit polyclonal anti-ADAM17 | Abcam | Cat# ab39162, RRID: AB_722565 |
| Rabbit polyclonal anti-14-3-3 epsilon | Cell Signaling Technology | Cat# 9635 RRID: AB_2217758 |
| Mouse monoclonal anti-GAPDH | Sigma-Aldrich | Cat# G8795 RRID: AB_1078991 |
| Mouse monoclonal anti- β-actin | Santa Cruz Biotechnology | Cat# sc-47778, RRID: AB_626632 |
| Rat monoclonal anti-HA-HRP, (clone 3F10) | Roche | Cat# 12013819001, RRID: AB_390917 |
| Goat polyclonal anti-rabbit-HRP | Bio-Rad | Cat# 170-6515, RRID: AB_11125142 |
| Horse polyclonal anti-mouse-HRP | Cell Signaling Technology | Cat# 7076, RRID: AB_330924 |
| Mouse monoclonal anti-beta-actin-HRP | Sigma-Aldrich | Cat# A3854, RRID: AB_262011 |
| **Bacterial and virus strains** |  |  |
| Stellar Competent Cells (an *E. coli* HST08 strain) | Takara Bio | Cat# 636763 |
| *E. coli* DH5α Competent Cells | GoldBio | Cat# CC-101-TR |
| *E. coli* DH10Bac Competent Cells | Thermo Fisher | Cat# 10361012 |
| **Chemicals, peptides, and recombinant proteins** |  |  |
| 1,10-Phenanthroline | Sigma-Aldrich | Cat# 131377-25G |
| cOmplete™, EDTA-free protease cocktail | Roche | Cat# 11873580001 |
| GW280264X (GW) | Generon | Cat# AOB3632-5 |
| DMEM high glucose medium | Sigma-Aldrich | Cat# D6429-500ML |
| Phenol Red-free OptiMEM | Gibco | Cat# 11058-021 |
| L-glutamine 200 mM (100x) | Gibco | Cat# 25030-024 |
| Penicillin-Streptomycin (P/S 100x) | Gibco | Cat# 15140-122 |
| Fetal Bovine Serum (FBS) | Sigma-Aldrich | Cat# F9665-500ML |
| FuGene HD Transfection Reagent | Promega | Cat# E2312 |
| PNPP tablet | Thermo Fisher Scientific | Cat# 34047 |
| Diethanolamine Substrate Buffer 5X concentrate | Thermo Fisher Scientific | Cat# 34064 |
| Bortezomib (BTZ) | Selleck Chemical | Cat# S1013 |
| Bafilomycin A1 | Santa Cruz Biotechnology | Cat# sc-201550A |
| ESF 921 Insect Cell Culture Medium | Expression Systems | Cat# 96-001-01 |
| TransIT-Insect transfection reagent | Mirus | Cat# MIR 6100 |
| Freestyle 293 expression medium | Gibco | Cat# 12338018 |
| Sodium butyrate | Millipore-Sigma | Cat# 303410 |
| Protease Inhibitor Cocktail | Medchemexpress | Cat# HY-K0010 |
| Benzonase nuclease | Millipore-Sigma | Cat# E1014-25KU |
| n-Dodecyl-β-D-Maltopyranoside (DDM) | Anatrace | Cat# D310 |
| Cholesteryl Hemisuccinate Tris Salt (CHS) | Anatrace | Cat# CH210 |
| Digitonin | Thermo Scientific | Cat# 407560050 |
| TCEP solution | Thermo Scientific | Cat# 77720 |
| **Deposited data** |  |  |
| iRhom1/ADAM17 complex | This paper | PDB: 12VV; EMDB: EMD-76788 |
| **Experimental models: Cell lines** |  |  |
| HEK293T cells | Laboratory of Matthew Freeman | RRID:CVCL_0063 |
| HEK293T iRhom1/iRhom2 double-knockout (DKO) | Künzel et al., 2018<sup>20</sup> | N/A |
| Sf9 insect cell | ATCC | Cat# CRL-1711; RRID:CVCL_0549 |
| HEK-293S GnTI- | ATCC | Cat# CRL-3022; RRID:CVCL_A785 |
| **Recombinant DNA** |  |  |
| pcDNA3.1 | Thermo Fisher Scientific | Cat# V790-20 |
| pcDNA3.1_hiR2_iso2_WT | This paper | N/A |
| pcDNA3.1_hiR1_WT | This paper | N/A |
| pcDNA3.1_hiR1_N458A/W460A | This paper | N/A |
| pcDNA3.1_hiR1_H668A | This paper | N/A |
| pcDNA3.1_hiR1_F675A | This paper | N/A |
| pcDNA3.1_hiR1_W766A/I767A | This paper | N/A |
| pcDNA3.1_hiR2_iso2_F647A | This paper | N/A |
| pcDNA3.1_hiR2_iso2_F401A | This paper | N/A |
| pcDNA3.1_hiR2_iso2_W738A | This paper | N/A |
| pcDNA3.1_hiR1_G462R | This paper | N/A |
| pcDNA3.1_hiR1_A538V | This paper | N/A |
| pcDNA3.1_hiR1_G665W | This paper | N/A |
| pLEX.puro-human iRhom1WT-3xHA | Künzel et al.<sup>20</sup> | N/A |
| pLEX.puro-human iRhom2WT-3xHA isoform1 | Künzel et al.<sup>20</sup> | N/A |
| pLEX.puro-human iRhom2WT-3xHA isoform2 | This paper | N/A |
| **Software and algorithms** |  |  |
| Prism 10 | GraphPad | https://www.graphpad.com/features |
| Fiji | Schindelin et al., 2012<sup>31</sup> | https://fiji.sc/ |
| FlowJo™ 10 | Becton, Dickinson and Company | https://flowjo.com/ |
| MotionCor2 | Zheng et al., 2017<sup>32</sup> | https://emcore.ucsf.edu/ucsf-motioncor2 |
| cryoSPARC v3.3.1 | Punjani et al., 2017<sup>33</sup> | https://cryosparc.com/ |
| RELION 3.1 | Zivanov et al., 2018<sup>34</sup> | https://www3.mrc-lmb.cam.ac.uk/relion/ |
| Coot 0.9.6 | Emsley et al., 2010<sup>35</sup> | https://www2.mrc-lmb.cam.ac.uk/personal/pemsley/coot/ |
| Phenix 1.16 | Adams et al., 2010<sup>36</sup> | http://www.phenix-online.org/ |
| PyMOL 2.3.4 | Schrödinger | https://pymol.org/2/ |
| MolProbity | Williams et al., 2018<sup>37</sup> | http://molprobity.biochem.duke.edu |
| UCSF Chimera 1.13 | Pettersen et al., 2004<sup>38</sup> | http://www.cgl.ucsf.edu/chimera |
| ChimeraX | Goddard et al., 2018<sup>39</sup> | https://www.cgl.ucsf.edu/chimerax/ |
| **Other** |  |  |
| Pierce™ anti-HA magnetic beads | Thermo Fisher Scientific | Cat# 88837 |
| Concanavalin A sepharose | Sigma-Aldrich | Cat# C9017-25ML |

### Bacterial strains

Stellar competent cells (an E. coli HST08 strain) or DH5α competent cells were used for molecular cloning and amplification of the recombinant plasmids. Bacteria were grown on LB broth with appropriate antibiotics.

### Cell lines and culture

Human embryonic kidney (HEK) 293T cells, iRhom1/iRhom2 double knockout (DKO) HEK 293T cells, HT-29 cells, HT-29 cells stably expressing UNC93B, iRhom1 and iRhom2 isoform1 and iRhom2 isoform2 were cultured in DMEM (Sigma-Aldrich) supplemented with 10% fetal bovine serum (FBS, Sigma-Aldrich), 2 mM L-Glutamine,100 U/ml penicillin and 100 μg/ml streptomycin (all Gibco). Cells were cultured at 37°C with 5% CO2 in a humidified cell culture incubator and were split twice a week using TrypLE Express (Gibco). Sf9 cells were cultured in ESF 921 Insect Cell Culture Medium at 27°C. HEK293S GnTI-cells were cultured in Freestyle 293 expression medium at 37°C.

### Molecular Cloning

The complementary DNA encoding human iRhom1, ADAM17, or FRMD8 was individually cloned into the pEG BacMam vector^40^. For full-length ADAM17, the coding sequence is followed by a TEV protease cleavage site and a C-terminal mScarlet tag. For iRhom1, a N-terminal mVenus tag and a 3C protease cleavage site are inserted before the coding sequence^41^. The expression cassettes containing individual genes were amplified and assembled into the pBIG1a vector using biGBac method^42^. The multigene expression construct containing iRhom1/ADAM17/FRMD8 was used for large-scale protein expression for cryo-EM.

For functional studies, wildtype iRhom1 cDNA was amplified from the previously published pLEX.puro-human iRhom1WT-3xHA construct ^20^ by PCR and was subcloned into pcDNA3.1(+) using In-Fusion HD Cloning Kit (Takara Bio, 639649). Mutant iRhom1 construct was generated by site-directed mutagenesis using the Cloned Pfu DNA polymerase AD (Agilent) according to the manufacturer’s instructions. Single colonies were picked and the sequences were verified by Sanger sequencing (Source BioScience, UK).

### Protein expression and purification for cryo-EM

The iRhom1/ADAM17 complex was expressed in HEK293S GnTI-cells. Baculoviruses were produced by transfecting Sf9 cells with bacmids generated from DH10Bac competent cells using TransIT (Mirus). After one or two rounds of amplification, viruses were used for cell transduction. When HEK293S GnTI− suspension cultures grown at 37 °C reached a density of ∼3.5 × 106 cells/ml, baculoviruses (10 % v/v) were added to initiate transduction. After 10–12 hrs, 10 mM sodium butyrate was supplemented to the cultures, and the culture temperature was shifted to 25 °C. Cells were collected at 60 hr post-transduction.

The cell pellet was resuspended using hypotonic buffer (10 mM NaCl, 1 mM MgCl2, 20 mM Tris pH 8, 2 mg/ml iodoacetamide, 0.1 mM TCEP, benzonase, and protease inhibitors) for 20 min. The cell lysate was then spun at 39,800g for 30 mins to sediment crude membranes. The membrane pellet was mechanically homogenized and solubilized in extraction buffer (20 mM DDM, 4 mM CHS, 150 mM NaCl, 20 mM Tris pH 8, 2 mg/ml iodoacetamide, 0.1 mM TCEP, benzonase, and protease inhibitors) for 1.5 h. Solubilized membranes were clarified by centrifugation at 39,800g for 45 mins. The supernatant was applied to the GFP nanobody-coupled glyoxal agarose resin, which was subsequently washed with 10 column volumes of wash buffer A (0.05 % digitonin, 150 mM NaCl, 0.1 mM TCEP, 4 mM NaATP, 4 mM MgCl2 and 20 mM Tris pH 8), followed by 7 column volumes of wash buffer B (0.05 % digitonin, 150 mM NaCl, 0.1 mM TCEP and 20 mM Tris pH 8). The washed resin was incubated with 3C protease overnight at a target protein to protease ratio of 40:1 (w/w) to cleave off mVenus and release the protein from the resin. The protein was eluted with wash buffer B, concentrated, and further purified by gel-filtration chromatography using a Superose 6 increase column equilibrated with SEC buffer (0.05 % digitonin, 150 mM NaCl, 0.1 mM TCEP, and 20 mM Tris pH 8). Peak fractions were pooled and concentrated for cryo-EM experiments or peptide cleavage assay.

### Cryo-EM sample preparation and data acquisition

Protein samples were concentrated to ∼7 mg/ml. Aliquots of 3.5 μl protein samples were applied to plasma-cleaned Quantifoil UltrAuFoil R1.2/1.3 300 mesh grids. After 25 s, the grids were blotted for 3 s and plunged into liquid ethane using a Vitrobot Mark IV (FEI) operated at 10 °C and 100% humidity. The grids were loaded onto a 300 kV Titan Krios transmission electron microscope for data collection. Raw movie stacks were recorded with a K3 camera at a pixel size of 0.649 Å on Krios. The nominal defocus range was 0.6–1.6 μm. Each micrograph was recorded with a total dose of 62.3 e−/Å and fractionated into 60 frames. Image acquisition parameters are summarized in Table S1.

### Cryo-EM Image Processing

The image stacks were firstly gain-normalized and corrected for beam-induced motion using MotionCor2^32^. Defocus parameters were estimated from motion-corrected images using cryoSPARC3^33^. Micrographs not suitable for further analysis were removed by manual inspection. Particle picking (blob picker and template picker) and 2D classifications were done in cryoSPARC3 (Figure S1A). After 2-3 rounds of 2D classifications, selected particles were used for iterative 3D classifications including ab initio reconstructions and heterogeneous refinements to remove suboptimal particles. The best classes were then subjected to nonuniform refinements for 3D reconstructions^43^. The refined particles were subjected to Bayesian polishing in RELION 3.1^34^. The polished particles were imported into cryoSPARC3 where additional nonuniform refinements were performed. The mask-corrected FSC curves were calculated in cryoSPARC3, and reported resolutions are based on the 0.143 criterion (Figure S1B and S1C). Local resolution estimations were performed in cryoSPARC3 (Figure S1D and S1E).

### Model building and Refinement

A model of iRhom1 was generated by AlphaFold^44^ and docked into the density map using Chimera^38^. A previously published ADAM17 structure (PDB code: 8SNL) was also fitted into cryo-EM density. The resulting complex model was then iteratively refined in Coot^35^ and Phenix^36^. Model validation was performed using Phenix and MolProbity^37^. Figures were prepared using PyMOL, Chimera, and ChimeraX^39^.

### DNA transfection

For HEK293T cells and iRhom1/iRhom2 DKO HEK293T cells, FuGENE® HD (Promega) was used, with a 4:1 ratio of transfection reagent (μl): DNA (μg), both of which were diluted in OptiMEM (Gibco).

### Lentiviral transduction of cell lines

HT-29 cells stably expressing iRhom1 and iRhom2 isoform1/2 were generated by lentiviral transduction using the pLEX.puro constructs as previously described ^9^. Cells were selected by adding 1 μg/ml puromycin.

### Flow cytometry with Cell surface labelling

All procedures were done on ice. Cells were washed with ice-cold PBS for 3 times and dissociated in FACS buffer (1% FBS in PBS). Cells were spun down at 400g for 4 mins and resuspend cells in 50ul FACS buffer containing primary antibodies, followed by incubation for 45mins. Cells were washed three times with FACS buffer and then resuspend cells in 50ul FACS buffer containing secondary antibodies, followed by incubation for 30 mins, protected from lights. Cells were then washed three times and were subjected to analysis on Cytoflex LX machine. Data were analysed using FlowJo.

### Co- Immunoprecipitation & Western Blotting analysis

Cells were washed with ice-cold PBS and lysed in Triton X-100 lysis buffer (1% Triton X-100, 150 mM NaCl, 50 mM Tris-HCl, pH 7.5) supplemented with EDTA-free complete protease inhibitor mix (Roche) and 10 mM 1,10-phenanthroline (Sigma-Aldrich). Lysates were cleared by centrifugation at 21,130 × g at 4°C for 15 minutes, and the supernatant was incubated with pre-washed anti-HA magnetic beads (Thermo Scientific) on a rotator at 4°C overnight. Beads were washed three times with Triton X-100 lysis buffer, and bound proteins were eluted using 2× SDS sample buffer (0.25 M Tris-HCl, pH 6.8, 10% SDS, 50% glycerol, 0.02% bromophenol blue) supplemented with 100 mM DTT. Samples were incubated at 65°C for 10 minutes before western blot analysis.

Tris-Glycine running buffer (25 mM Tris, 192 mM glycine, 0.1% SDS) was used for Novex 8-16% Tris-Glycine Mini Gels with WedgeWell format (Thermo Scientific). MOPS running buffer (50 mM MOPS, 50 mM Tris, 0.1% SDS, 1 mM EDTA) was used for Bis-Tris gels. Proteins were transferred to a methanol-activated polyvinylidene difluoride (PVDF) membrane (Millipore) using Bis-Tris or Tris-Glycine transfer buffers. Membranes were blocked and incubated with primary and secondary antibodies in 5% milk in PBST (0.1% Tween 20). After incubation with secondary antibodies at room temperature for 1 hour, membranes were washed with PBST.

### Concanavalin A (ConA) Enrichment

Cleared cell lysates (supernatants) were incubated with 20 μl concanavalin A sepharose (Sigma-Aldrich, C9017-25ML) on a rotator at 4°C overnight. Beads were pelleted by centrifugation at 1500 × g for 2 minutes at 4°C and washed five times with Triton X-100 lysis buffer. Glycoproteins were eluted using 2 × LDS buffer (Invitrogen) supplemented with 25% sucrose and 50 mM DTT, followed by incubation at 65°C for 10 minutes. Samples were then analysed by western blot using 4–12% Bis-Tris NuPAGE gradient gels (Invitrogen).

### Alkaline phosphatase (AP) - Shedding assay

Cells were seeded in poly-L-lysine (PLL, Sigma-Aldrich)-coated 24-well plates in triplicate 24 hours before transfection. Cells were transfected with 200ng iRhom1 variants and 150 ng of alkaline phosphatase (AP) - conjugated substrates using FuGENE HD (Promega, E2312). 24 hours post-transfection, cells were washed with PBS and incubated for 20 hours in 300 μl Phenol Red-free OptiMEM (Gibco, 11058-021). Culture medium (supernatant) was then collected, and cells were lysed in 300 μl Triton X-100 lysis buffer supplemented with EDTA-free protease inhibitors (Roche) and 10 mM 1,10-phenanthroline (Sigma-Aldrich, 131377–5G). 100 μl of supernatant and 100 μl of diluted cell lysate were independently incubated with 100 μl AP substrate p-nitrophenyl phosphate (PNPP; Thermo Fisher Scientific, 37620) at room temperature. Absorbance was measured at 405 nm using a SpectraMax M3 plate reader (Molecular Devices). Substrate release was quantified as the ratio of supernatant signal to total signal (supernatant + cell lysate).

## Acknowledgement

We thank Professor Christoph Tang (Dunn School, Oxford, UK) for generously sharing the HT-29 cells and acknowledge the cells lines (HT-29 + Unc93B, HT-29 + WT iRhom1) were generated by Dr Colin Adrain during his postdoc time in the Freeman lab. This work was supported by NIH (R01GM143282 and R01NS133147) and ALSAC to C.-H.L.; Wellcome Trust Investigator awards to M.F. (101035/Z/13/Z and 220887/Z/20/Z).

## Author contributions

F.L. designed, performed, and analyzed cellular functional experiments. H.Z. designed, performed, and analyzed structural experiments. Y.D. assisted in structural analysis. C.-H.L. and M.F. conceived the research and supervised the project. F.L., C.-H.L., and M.F. wrote the manuscript. All authors contributed to manuscript preparation.

## Declaration of interests

The authors declare no competing financial interests.

## Notes

### Competing Interest Statement

The authors have declared no competing interest.

