## Supplemental information for "A sterol-binding pocket in iRhom1 underlies paralog-specific regulation of the sheddase ADAM17"

| Structure | ADAM17/iRhom1 |
|---|---|
| PDB | 12VV |
| EMDB | EMD-76788 |
| **Data collection/ processing** | |
| Magnification | 130,000x |
| Voltage (kV) | 300 |
| Pixel size (Å) | 0.649 |
| Defocus range (μm) | 0.6-1.6 |
| Electron exposure ($e^-/Å^2$) | 62.3 |
| Symmetry imposed | C1 |
| Initial particles (No.) | ~9.5 million |
| Final particles (No.) | 335,665 |
| Map resolution (Å) | 2.50 |
| FSC threshold | 0.143 |
| Map resolution range (Å) | 40-2.2 |
| **Refinement** | |
| Model Resolution (Å) | 2.7 |
| FSC threshold | 0.5 |
| Map sharpening B-factor ($Å^2$) | -72.8 |
| Model composition | |
| Non-hydrogen atoms | 9065 |
| Protein residues | 1139 |
| Ligand | 3 |
| *B*-factors ($Å^2$) | |
| Protein | 47.92 |
| Ligand | 38.26 |
| R.m.s. deviations | |
| Bond lengths (Å) | 0.005 |
| Bond angles (°) | 1.030 |
| Validation | |
| MolProbity score | 1.74 |
| Clashscore | 6.85 |
| Rotamers outliers (%) | 0.00 |
| Ramachandran plot (%) | |
| Favored | 94.79 |
| Allowed | 5.21 |
| Outliers | 0.00 |

**Table S1.** Cryo-EM data collection, processing, and refinement statistics

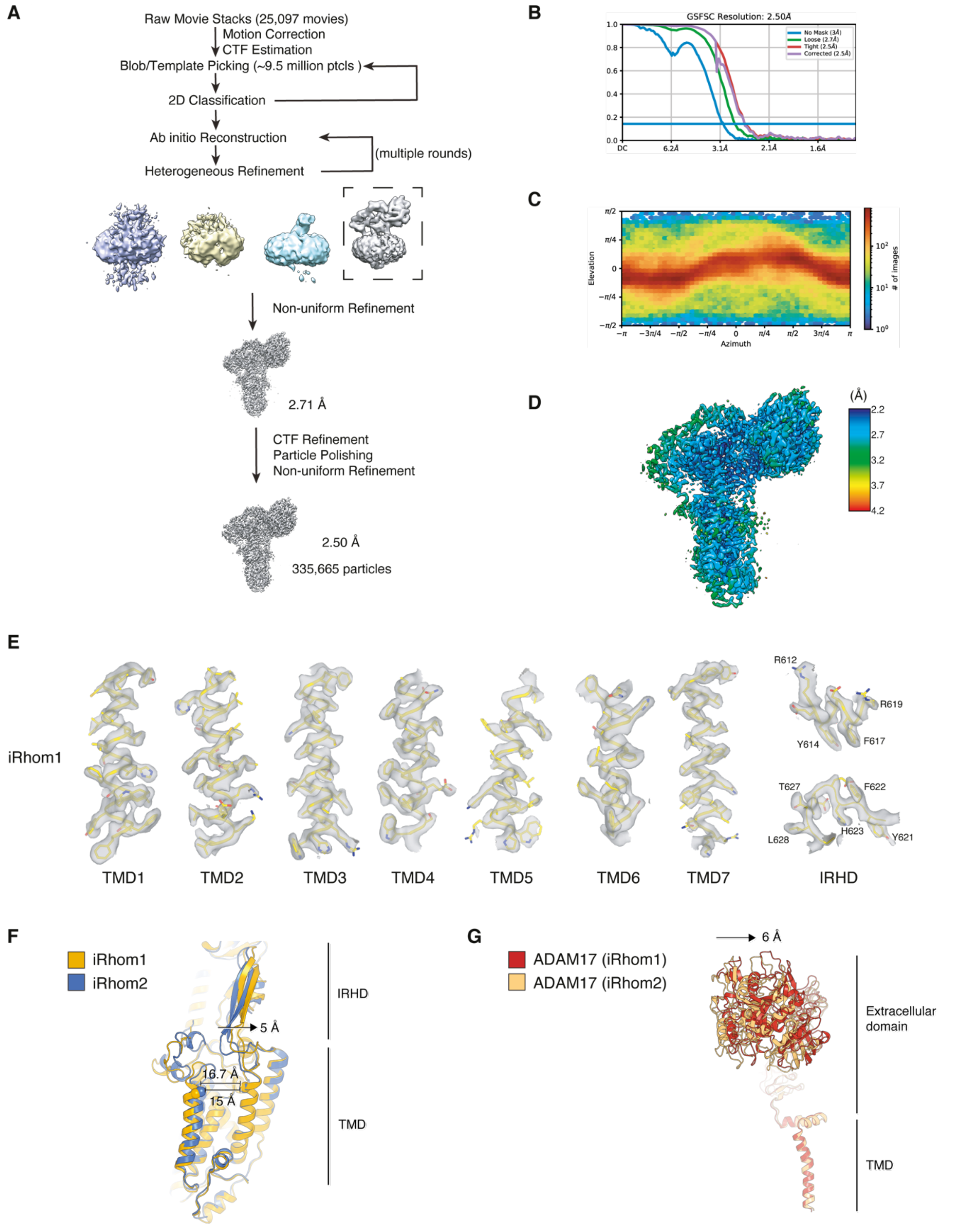


**Figure S1. Cryo-EM processing, validation and structural comparison of the iRhom1/ADAM17 complex.** (A) Cryo-EM data-processing workflow for the human iRhom1/ADAM17 complex. (B) Gold-standard Fourier shell correlation curve for the final iRhom1/ADAM17 reconstruction. (C) Angular distribution of particles for the final

3D reconstructions. (D) Local-resolution map of the iRhom1/ADAM17 complex, coloured according to local resolution. (E) Representative cryo-EM densities for iRhom1 transmembrane helices and the IRHD, showing the quality of the map in the modelled regions. (F) Structural superposition of the iRhom1 and iRhom2 transmembrane regions. iRhom1 and iRhom2 are shown in yellow and blue, respectively. Differences in the relative positions of the IRHD and transmembrane core are indicated. (G) Structural comparison of ADAM17 in the iRhom1/ADAM17 and iRhom2/ADAM17 complexes, showing a displacement of the ADAM17 extracellular domain between the two complexes.

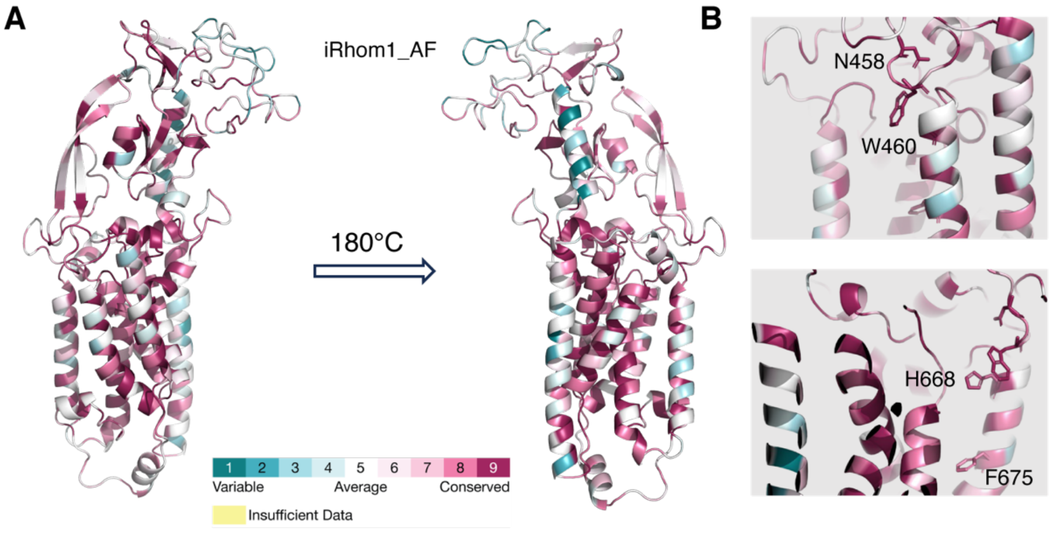


**Figure S2. Conservation analysis of the iRhom1 sterol-binding pocket.** (A) Human iRhom1 coloured according to the residue conservation analysed by ConSurf. AlphaFold model was used to show all residues of TMD and extracellular domain. (B) Close-up views of the iRhom1 sterol-binding pocket showing conservation score of residues N458, W460, H668 and F675 (side chains were shown as sticks).

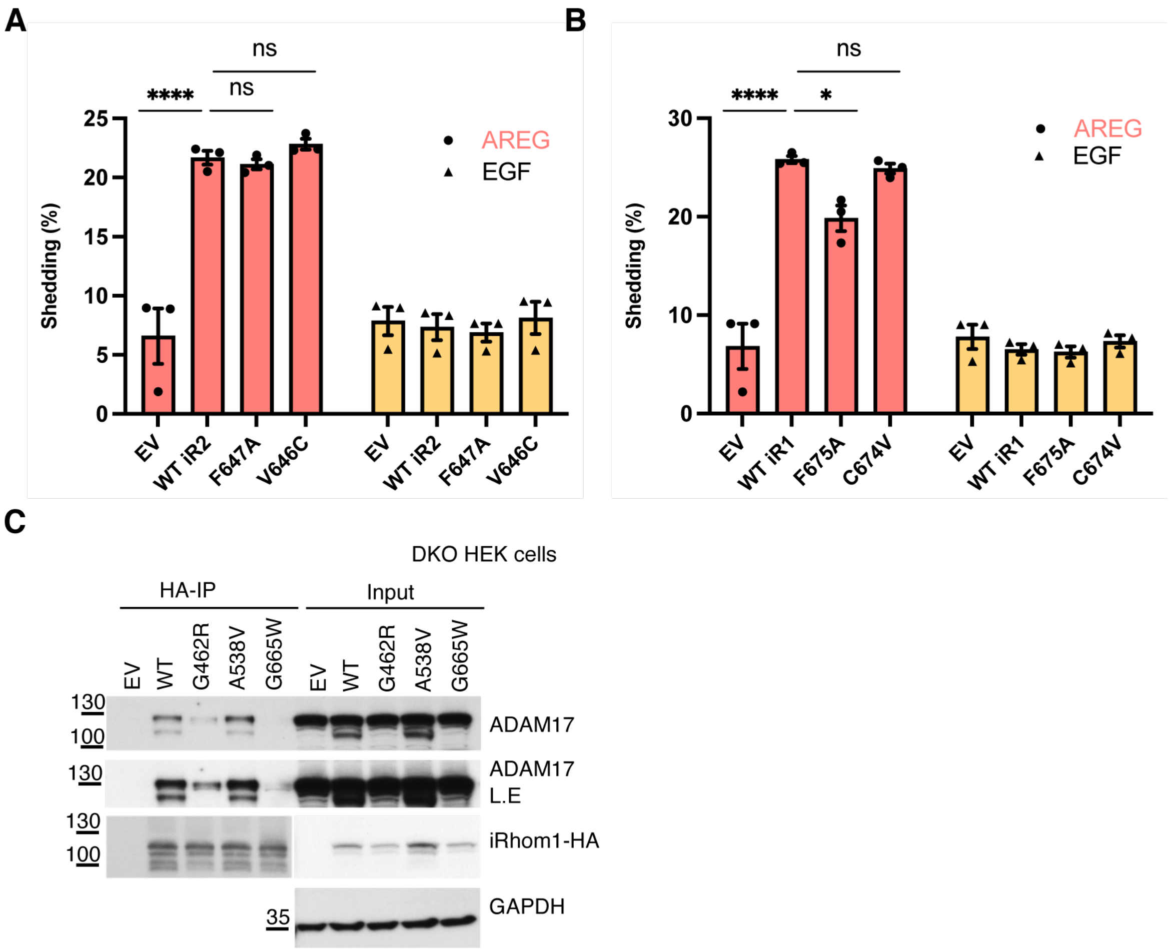


**Figure S3. Additional functional analysis of iRhom1 and iRhom2 sterol-pocket and disease-associated variants.** (A) ADAM17-dependent shedding assay in iRhom1/2 DKO cells expressing empty vector, wild-type iRhom2 isoform 2, or the indicated iRhom2 variants. Shedding of alkaline phosphatase-tagged AREG and EGF substrates was quantified. Data are shown as mean ± SD; ns = not significant, **** = $p < 0.0001$. (n=3, independent experiment). (B) ADAM17-dependent shedding assay in iRhom1/2 DKO cells expressing empty vector, wild-type iRhom1, or the indicated iRhom1 variants. Shedding of alkaline phosphatase-tagged AREG and EGF substrates was quantified. Data are shown as mean ± SEM; ns = not significant, * = $p < 0.05$, **** = $p < 0.0001$. (C) HA immunoprecipitation analysis in iRhom1/2 double-knockout HEK293T cells expressing empty vector, wild-type iRhom1, the control A538V variant, or the disease-associated G462R and G665W variants. Co-precipitated ADAM17 and iRhom1-HA were detected by immunoblotting; GAPDH served as a loading control.
